# Patterns of microbiome variation among infrapopulations of permanent bloodsucking parasites

**DOI:** 10.1101/2020.05.27.118331

**Authors:** Jorge Doña, Stephany Virrueta Herrera, Tommi Nyman, Mervi Kunnasranta, Kevin P. Johnson

**Affiliations:** Illinois Natural History Survey, Prairie Research Institute, University of Illinois at Urbana-Champaign, 1816 S. Oak St., Champaign, Illinois 61820, USA; Departamento de Biología Animal, Universidad de Granada, Granada, 18001, Spain; Department of Ecosystems in the Barents Region, Norwegian Institute of Bioeconomy Research, Svanhovd 35, 9925 Svanvik, Norway; Department of Environmental and Biological Sciences, University of Eastern Finland, Yliopistokatu 7, 80101 Joensuu, Finland; Natural Resources Institute Finland, Joensuu, Finland

**Keywords:** genome-resolved metagenomics, host-symbiont, intraspecific variation, lice, microbiota, shotgun metagenomics, symbiont

## Abstract

While interspecific variation in microbiome composition can often be readily explained by factors such as host species identity, there is still limited knowledge of how microbiomes vary at scales lower than the species level (e.g., between individuals or populations). Here, we evaluated variation in microbiome composition of individual parasites among infrapopulations (i.e., populations of parasites of the same species living on a single host individual). To address this question, we used genome-resolved and shotgun metagenomic data of 17 infrapopulations (balanced design) of the permanent, bloodsucking seal louse *Echinophthirius horridus* sampled from individual Saimaa ringed seals *Pusa hispida saimensis*. Both genome-resolved and read-based metagenomic classification approaches consistently show that parasite infrapopulation identity is a significant factor that explains both qualitative and quantitative patterns of microbiome variation at the intraspecific level. This study contributes to the general understanding of the factors driving patterns of intraspecific variation in microbiome composition, especially of bloodsucking parasites, and has implications for understanding how well-known processes occurring at higher taxonomic levels, such as phylosymbiosis, might arise in these systems.

## Introduction

Patterns of inter- and intraspecific variation in microbiome composition of animals have received much attention because the microbiome may influence many biological processes that have considerable effects on the host (Clemente et al. 2012; Le Chatelier et al. 2013; Rothschild et al. 2018; Rudman et al. 2019; Velazquez et al. 2019). For instance, particular microbiome compositions have been found to drive genomic adaptation (Rudman et al. 2019) or to confer protection against pathogens (Velazquez et al. 2019).

In general, both stochastic (e.g., dispersal, or ecological drift) and deterministic (e.g., host immunological regulation, or microbe–microbe interactions) processes operate across multiple spatial scales to shape the composition of animal microbiomes (Adair and Douglas 2017; Kohl 2020). In particular, among the many determinants shaping microbiome composition, host species identity has been repeatedly identified as a key factor determining the composition of animal microbiomes (Brooks et al. 2016; Mazel et al. 2018; Nishida and Ochman 2018; Lutz et al. 2019; Knowles et al. 2019; Lim and Bordenstein 2020; Song et al. 2020). In other words, microbiomes of individuals of the same species tend to be more similar than to those of another species. This pattern is generally the result of filtering microbial taxa by the host (e.g., through host diet, habitat, or immune system, Adair and Douglas 2017) or result from host-microbe coevolution (Lim and Bordenstein 2020). When this process exhibits phylogenetic signal, the pattern is known as phylosymbiosis (i.e., microbial community relationships that recapitulate the phylogeny of their host, Brucker and Bordenstein 2013; Brooks et al. 2016; Lim and Bordenstein 2020). Nonetheless, several aspects of the variation of animal microbiomes are yet to be better understood (Lim and Bordenstein 2020). In particular, for non-human animals, there is still much to learn about how microbiomes vary at scales below the species level, such as between populations (Blekhman et al. 2015; Kohl et al. 2018; Rothschild et al. 2018; Campbell et al. 2020; Fountain-Jones et al. 2020) or ecotypes (Agany et al. 2020).

An area of focus on understanding intraspecific variation in microbiome composition has been bloodsucking parasites. In these parasites, previous studies have consistently found a major role of the host species in shaping microbiome composition in the parasites (Osei-Poku et al. 2012; Zhang et al. 2014; Swei and Kwan 2017; Zolnik et al. 2018; Landesman et al. 2019; Lee et al. 2019; Muturi et al. 2019). However, in ticks (*Ixodes scapularis*), host individual identity of the blood meal was even more important than host species identity in explaining microbiome composition (Landesman et al. 2019). These results suggested that individual host identity of the blood meal might be an important factor that shapes parasite microbiomes at the intraspecific level (Landesman et al. 2019). In theory, microbiomes of individual bloodsucking parasites could vary due to: 1) the individual parasite immune system that may impose selection on different bacterial taxa (Blekhman et al. 2015; Suzuki et al. 2019), 2) differences in the source of the blood meal that may transfer or disperse particular bacterial taxa, or modulate bacteria by creating specific conditions during digestion (Rothschild et al. 2018), 3) microbe–microbe interactions (Hassani et al. 2018), and 4) stochastic processes (e.g., ecological drift) (Lankau et al. 2012). However, for most species, and for bloodsucking parasites in particular, the nature of intraspecific variation in microbiomes and the relative importance of factors shaping this variation remain understudied.

Sucking lice (Phthiraptera: Anoplura) are permanent blood-feeding ectoparasites that live in the fur or hairs of mammals. The sucking lice of pinnipeds (seals, sea lions, and walrus) are of particular interest because of their need to adapt to the aquatic lifestyle of their hosts (Durden and Musser 1994; Leonardi et al. 2013). There is evidence that the sucking lice of seals and sea lions have codiversified with their hosts (Kim 1971, 1975, 1985; Leonardi et al. 2019). In addition, the sucking lice of pinnipeds represent an interesting system in which to study the variation in microbiome composition and the drivers of this variation at an intraspecific level because: 1) these lice have well defined, isolated populations (infrapopulations) on individual seal hosts, due to an expected low rate of horizontal dispersal among host individuals, which is only possible during the seals’ haul-out periods on land or ice (Kim 1985; Leonardi et al. 2013, 2019); and 2) these lice feed only upon the blood of their host (Snodgrass 1944; Kim 1985), so that it can be assumed that individuals from the same infrapopulation feed upon “exactly” the same resource (i.e., the blood of the individual seal on which they occur).

Here, we used genome-resolved approaches (the construction of draft microbial genomes from short-read shotgun sequencing data; Bowers et al. 2017; Uritskiy et al. 2018) and metagenomic classification tools (taxonomic classification of individual sequencing reads; Menzel et al. 2016) to infer patterns of microbiome variation among individuals of the sucking seal louse *Echinophthirius horridus* (von Olfers, 1816) inhabiting individual Saimaa ringed seals *Pusa hispida saimensis* (Nordquist, 1899). Our sampling design, involving analysis of two individual lice from each of 17 seals, allowed us to evaluate the degree to which variation in microbiome composition among individual lice is explained by the infrapopulation (the identity of the seal host).

## Materials and Methods

### Sampling, DNA extraction, and sequencing

Thirty-four individual lice were sampled from 17 individual Saimaa ringed seals (*Pusa hispida saimensis*), which is an endemic endangered landlocked subspecies of the ringed seal living in freshwater Lake Saimaa in Finland (e.g., Nyman et al. 2014). Individual lice were collected from seals found dead or from seals that were live-captured for telemetry studies (e.g., Niemi et al. 2019), and placed in 2-ml screw-cap tubes with 99.5% ethanol. Lice from a single seal individual were put in the same tube. Prior to DNA extraction, each louse individual was rinsed with 95% ethanol and placed alone in a new sterile vial; then, the remaining ethanol was evaporated at room temperature.

Whole lice were ground up individually, and genomic DNA was extracted using the Qiagen QIAamp DNA Micro Kit (Qiagen, Valencia, CA, U.S.A.). The standard protocol was modified so that specimens were incubated in ATL buffer and proteinase K at 55 (insert degree) C for 48 h instead of the recommended 1 – 3 h, as well as by substituting buffer AE with buffer EB (elution buffer). This was done to ensure maximal yield (greater than 5 ng) of DNA from each louse. Each DNA extract was quantified with a Qubit 2.0 Fluorometer (Invitrogen, Carlsbad, CA, U.S.A.) following the manufacturer’s recommended protocols.

Shotgun genomic libraries were prepared from the extracts with Hyper Library construction kits (Kapa Biosystems, Wilmington, MA, U.S.A.), and the libraries were quantitated by qPCR and 150 bp pair-end sequenced on one lane of an Illumina NovaSeq 6000 sequencer (Albany, New York). FASTQ files from sequence data were generated and demultiplexed with bcl2fastq v.2.20. All library preparations, sequencing, and FASTQ file generation were carried out at the Roy J. Carver Biotechnology Center (University of Illinois, Urbana, IL, U.S.A.). Raw reads were subsequently deposited to the NCBI GenBank SRA database (Table S1).

### Metagenomic analyses

For the genome-resolved metagenomic analyses, we used the metaWRAP v1.1.5 pipeline (Uritskiy et al. 2018) along with all the recommended databases (i.e., Checkm_DB, NCBI_nt, and NCBI_tax). We used the metaWRAP Read_qc module with default parameters to quality trim the reads and to de-contaminate each sample from host reads. For decontamination, we ran a de-novo genome assembly of an individual louse of the same species, not included in this study, and with a high amount of sequencing data (“Echor52”) in Abyss v2.0.1 (Jackman et al. 2017). Finally, we filtered out all non-bacterial reads from the contig file using Blobtools v1.0.1 (Laetsch and Blaxter 2017) and used this file to decontaminate all the other samples with the metaWRAP Read_qc module. Next, we co-assembled reads from all the samples with the metaWRAP Assembly module (--usemetaspades option) (Nurk et al. 2017). For this assembly, and because of memory limitations, we used BBNorm (sourceforge.net/projects/bbmap/) before assembly to reduce the coverage of the concatenated FASTQ file to a maximum of 100X and to discard reads with coverage under 3X. We binned reads with the metaWRAP Binning module (--maxbin2 --concoct --metabat2 options) (Alneberg et al. 2014; Wu et al. 2016; Kang et al. 2019) and then consolidated the resulting bins into a final bin set with both metaWRAP’s Bin_refinement module (-c 50 -x 10 options) and the Reassemble_bins module. We quantified the bins resulting from the Bin_refinement module with Salmon (Patro et al. 2017) using the Quant_bins module with default parameters. Finally, we classified bins using the Classify_bins module. This module uses Taxator-tk, which gives highly accurate but conservative classifications (Dröge et al. 2015). Accordingly, we also uploaded our final metagenome-assembled genomes (MAGs) to MiGA for a complementary analysis to determine the most likely taxonomic classification and novelty rank of the bin (Rodriguez-R et al. 2018). We used the NCBI Genome database (Prokaryotes; February 26, 2020 version) for this analysis.

For the metagenomic classification of reads, we used the metagenomic classifier Kaiju (Menzel et al. 2016) with Reference database: nr (Bacteria and Archaea; Database date: 2017-05-16). We used the default parameters for these analyses – SEG low complexity filter: yes; Run mode: greedy; Minimum match length: 11; Minimum match score: 75; Allowed mismatches: 5. We then converted Kaiju’s output files into a summary table at the genus and species level and filtered out taxa with low abundances (<0.1 % of the total reads). We also removed poorly identified taxa because they would artificially increase the similarity between our samples. Specifically, the following taxa were excluded: “NA”, “Viruses”, “archaeon”, “uncultured bacterium”, “uncultured Gammaproteobacteria bacterium” (Table S2 and S3).

Lastly, we used Decontam v1.2.1 to filter out bacterial taxa exhibiting known statistical properties of contaminants (Davis et al. 2018). We used the frequency method (*isContaminant* function) which is based on the inverse relationship between the relative abundance of contaminants and sample DNA concentration, and also has been found suitable for samples dominated by host DNA (Willner et al. 2012; Lusk 2014; Salter et al. 2014; Jervis-Bardy et al. 2015; Hooper et al. 2019; McArdle and Kaforou 2020). As input for Decontam analyses, we used the aforementioned total DNA concentration values. Then, as recommended, we explored the distribution of scores assigned by Decontam to assign the threshold according to bimodality between very low and high scores (Davis et al. 2018). For the MAGs matrix, no bimodality was found, and thus we used the 0.1 default value (Fig S1a). None of the MAGs were classified as contaminants, according to Decontam. For Kaiju matrices, a 0.3 threshold value was selected for the species matrix (Fig S1b) and 0.31 for the genus matrix (Fig S1c). Decontam filtered out a single species (*Clostridia* bacterium k32) from the species matrix and two genera (*Cupriavidus* and *Massilia*) from the genus matrix.

### Statistical analyses

To visualize similarities of microbiome composition among louse individuals from the same or different individual seal hosts, we constructed non-metric multidimensional scaling (NMDS) ordinations based on Bray–Curtis and Jaccard (binary = T) dissimilarities using the phyloseq v1.26-1 R package (McMurdie and Holmes 2013). For the genome-resolved metagenomic analyses, we used the normalized MAGs compositional matrices resulting from Salmon. Specifically, these MAG counts are standardized to the individual sample size (MAG copies per million reads) and thus allow between-sample comparisons. For the Kaiju analyses, we used the rarefy_even_depth function of phyloseq (without replacement as in the hypergeometric model) to rarefy samples to the smallest number of classified sequences per individual observed (85,513, and 71,267 reads in genus and species matrices, respectively) (Weiss et al. 2017). To assess the influence of individual host identity on the microbiome composition of louse individuals, we conducted a permutational multivariate analysis of variance (PERMANOVA) (Anderson and Walsh 2013; Anderson 2014). PERMANOVA analyses were done using the *adonis2* function in vegan v2.5–4 (Oksanen et al. 2019), based on Bray–Curtis and Jaccard distance matrices with 100 iterations. In PERMANOVA analyses, for the individual host identity factor, our within-group sample size (n=2) was smaller than both the total number of groups (n=17) and the total sample size (n=34). Thus, to account for a potential deviation in F-statistics and R^2^ values (Kelly et al. 2015), we wrote an R simulation that randomly subsampled the infrapopulations from which the louse came (5 infrapopulations per iteration). We ran 10 iterations and ran a PERMANOVA analysis for each iteration. Note that, for a few iterations, subsampled samples were too similar and PERMANOVA could not be done. In addition, we ran PERMANOVA analyses to explore additional factors (louse sex: male, female; sequencing lane: 1, 2; and host status: dead, alive) that may explain variance in microbiome composition. Furthermore, we included significant factors as the first factors of the host identity PERMANOVA models (i.e., to obtain the variance explained by host identity after accounting for the variance explained by that factor). We also restricted the groups in which permutations could be done to only those with the same value of that vector using the strata argument (e.g., for a sample collected from a dead host, and for the host-status factor, permutations could only be done among other dead hosts). Lastly, we ran a Mantel test using the *mantel* function in vegan (method=spearman, permutations=9999) to explore if host locality (i.e., the coordinates in which each host was sampled) correlated with the microbiome composition of louse individuals. For this analysis, we ran 10 iterations of an R simulation in which we randomly subsampled one louse sample for each individual host and ran a Mantel test for each iteration. The following packages were used to produce the plots: ggplot2 v3.1.0.9 (Wickham 2016), grid v3.5.3 (R Core Team 2019), gridExtra v.2.3 (Auguie 2016), ggrepel v0.8.0 (Slowikowski et al. 2019), ggpubr v.0.2.5 (Kassambara 2018), and ggsci v2.9 (Xiao 2018).

## Results

From the genome-resolved metagenomics pipeline, 13 high-quality bacterial metagenome-assembled genomes (MAGs) were obtained (Table 1; Fig 1). According to MiGA analyses, 10 of them (77%) likely belong to a species not represented in the NCBI Genome database.

**Figure 1.**
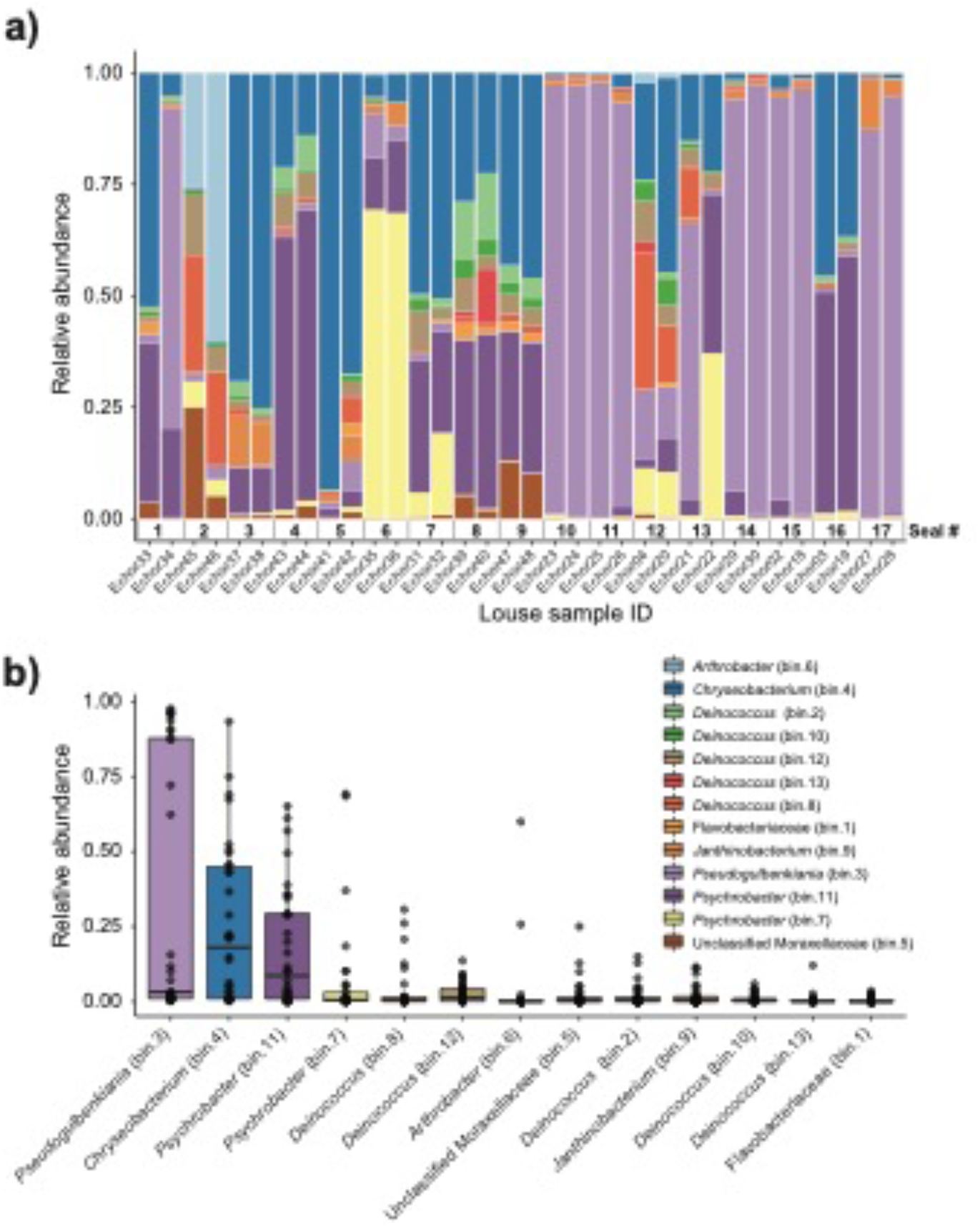
Genome-resolved metagenomic data. a) Stacked bar plot showing the relative abundances of MAGs in each louse sample. Note that samples are ordered according to host (i.e., samples from the same host are next to each other). b) Boxplot summarizing the relative abundance of each MAG across the louse samples.

**Table 1.**
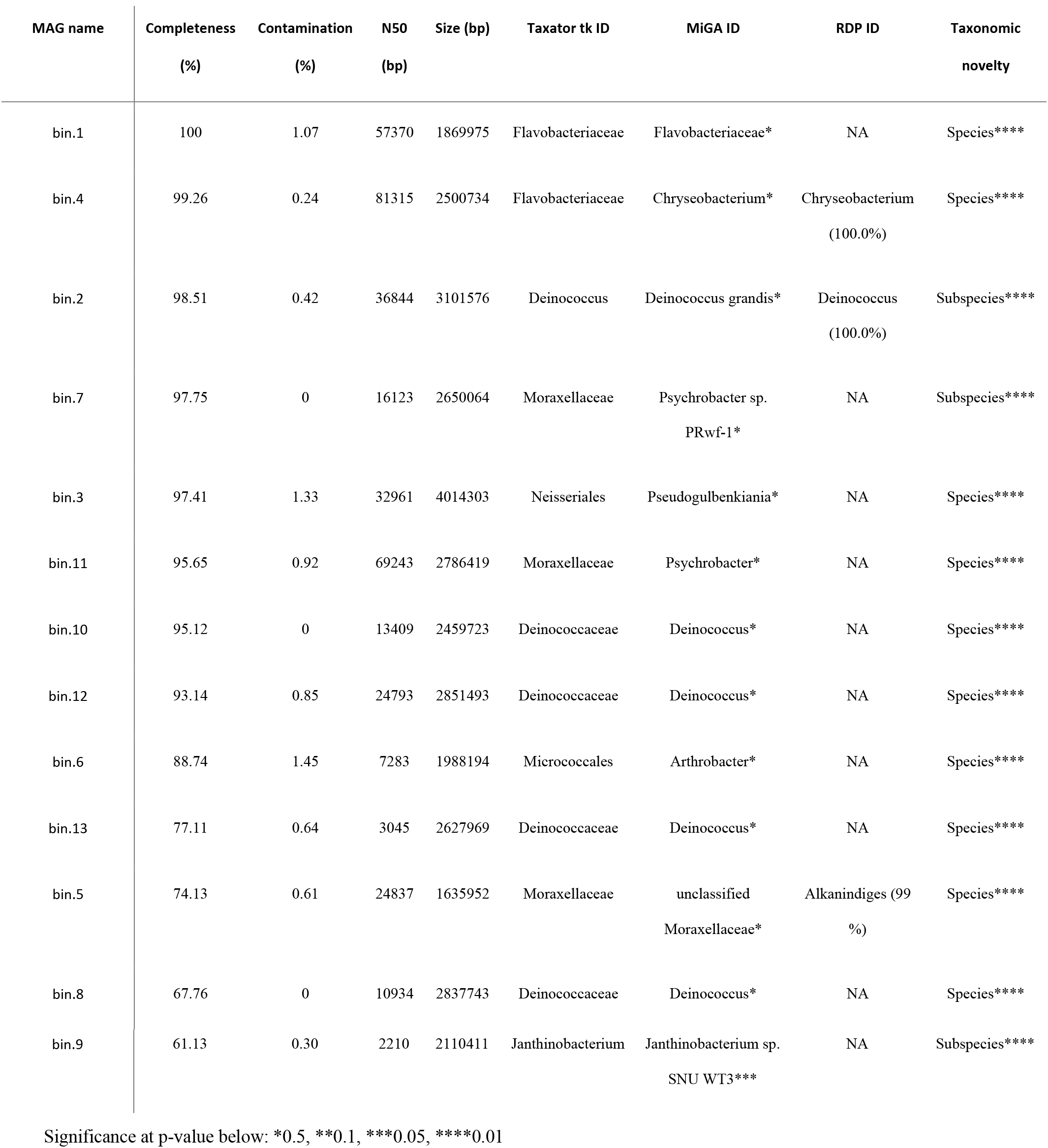
Statistics of the MAGs assembled. MAG name indicates the name given to that bin for this study (e.g., in Figure 1). The highest taxonomic rank with p-value ≤ 0.5 is shown in MiGA ID. RPD ID is the result of the identification analysis using rRNA genes (16S) implemented in MiGA; % indicates confidence in identification. Taxonomic novelty is a MiGA analysis that indicates the taxonomic rank at which the MAG represents a novel taxon with respect to the NCBI Genome database; highest taxonomic rank with p-value ≤ 0.01 are shown.

Kaiju analyses recovered a higher diversity of microorganisms than did the genome-resolved approach. These differences are likely because of the quality-filtering parameters used in the genome-resolved metagenomics pipeline (i.e., these taxa may have been discarded because the completeness values of their bins were lower than 50% and/or their contamination values were higher than 10%). Nevertheless, bacterial taxa found in the genome-resolved metagenomic approach were generally found also in Kaiju and with similar relative abundances (Fig 2), and a similar pattern was found also at the species level (Fig S2).

**Figure 2.**
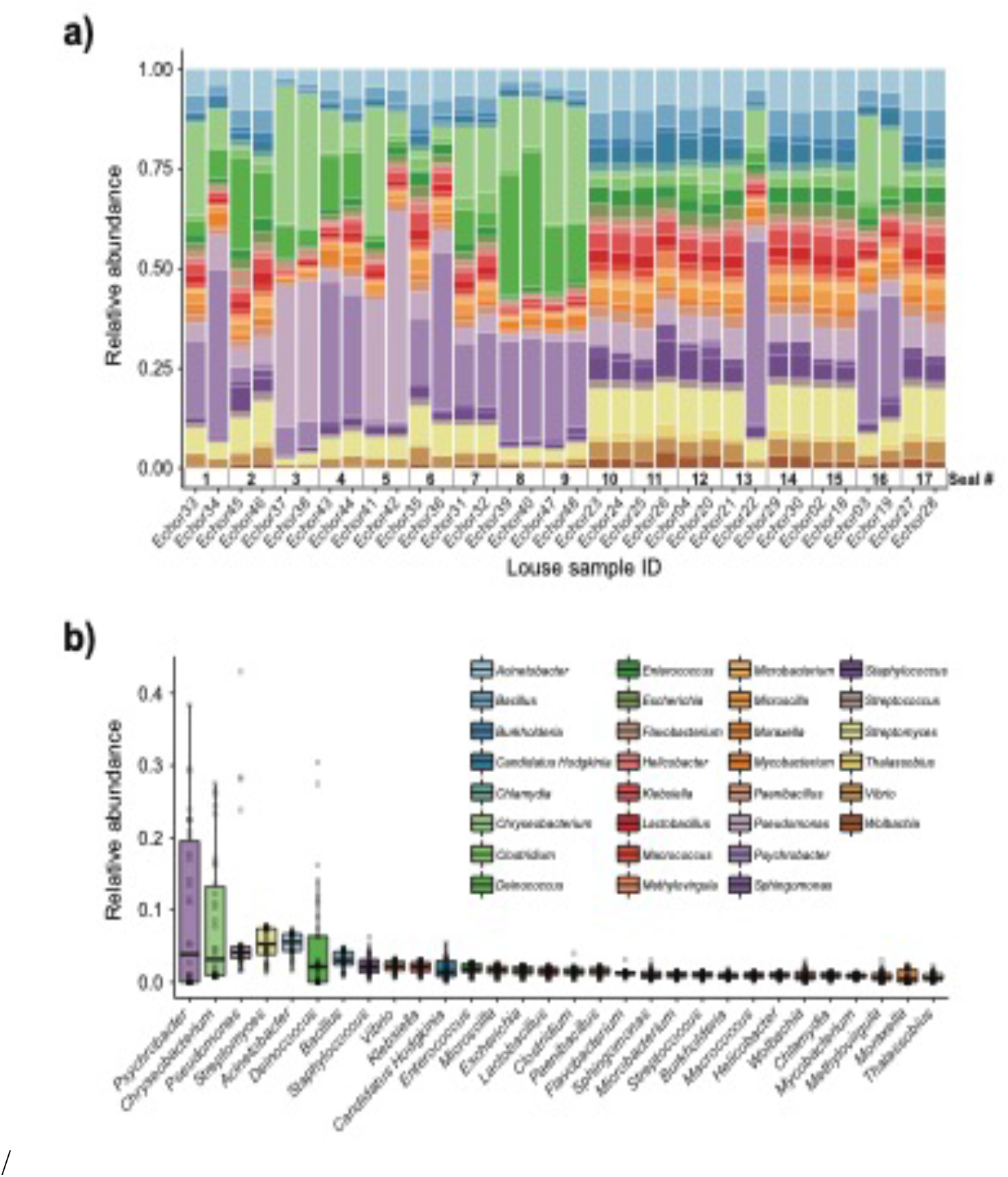
Kaiju data. a) Stacked bar plot showing bacterial relative abundances in each seal louse sample. Note that samples are sorted according to host individual (i.e., samples from the same host are next to each other). b) Boxplot summarizing the relative abundance of each taxon across all louse samples.

Ordination and PERMANOVA analyses show a major role of infrapopulation identity in explaining microbiome composition for both presence–absence and quantitative data. In the genome-resolved metagenomic dissimilarity matrices, most (>84% in all cases) of the variance was explained by infrapopulation identity (PERMANOVA: Bray-Curtis, R^2^= 0.857, F=6.419, P=0.001, Fig 3a; Jaccard, R^2^= 0.842, F= 5.671, P=0.001; Fig S3a). Results from the simulations were in line with the results of the regular model, and thus support that our results were not biased by the sampling design [PERMANOVA: Bray-Curtis, R^2^ (min= 0.65, max= 0.98, mean= 0.78); P (min= 0.001, max= 0.019, n<0.05= 10/10); Jaccard, R^2^ (min= 0.66, max= 1, mean= 0.86), P (min= 0.001, max= 0.106, n<0.05= 5/7)]. From all the additional factors examined, only host status (i.e., dead, alive) explained a significant amount of variance [PERMANOVA: Bray-Curtis, Host status: R^2^= 0.28, F= 12.72, P= 0.001, Louse sex: R^2^= 0.08, F= 0.9, P= 0.554, Sequencing lane: R^2^= 0.01, F= 0.38, P= 0.878; Jaccard, Host status: R^2^= 0.13, F= 4.93, P= 0.002, Louse sex: R^2^= 0.03, F= 0.28, P= 0.867, Sequencing lane: R^2^= 0, F= −0.01, P= 1; Mantel test, locality, Bray-Curtis: ρ (min= −0.09, max= −0.09, mean= −0.09), P (min= 0.8749, max= 0.8867, n<0.05= 0/10); Jaccard: ρ (min= −0.29, max= −0.29, mean= −0.29), P (min= 0.97742, max= 0.9777, n<0.05= 0/10)]. Including host status in PERMANOVA analyses did not alter the results on the major influence of host identity in explaining microbiome composition (PERMANOVA: Host identity, Bray-Curtis, R^2^= 0.57, F= 4.58, P= 0.001; Jaccard, R^2^= 0.71, F= 5.09, P= 0.002).

**Figure 3.**
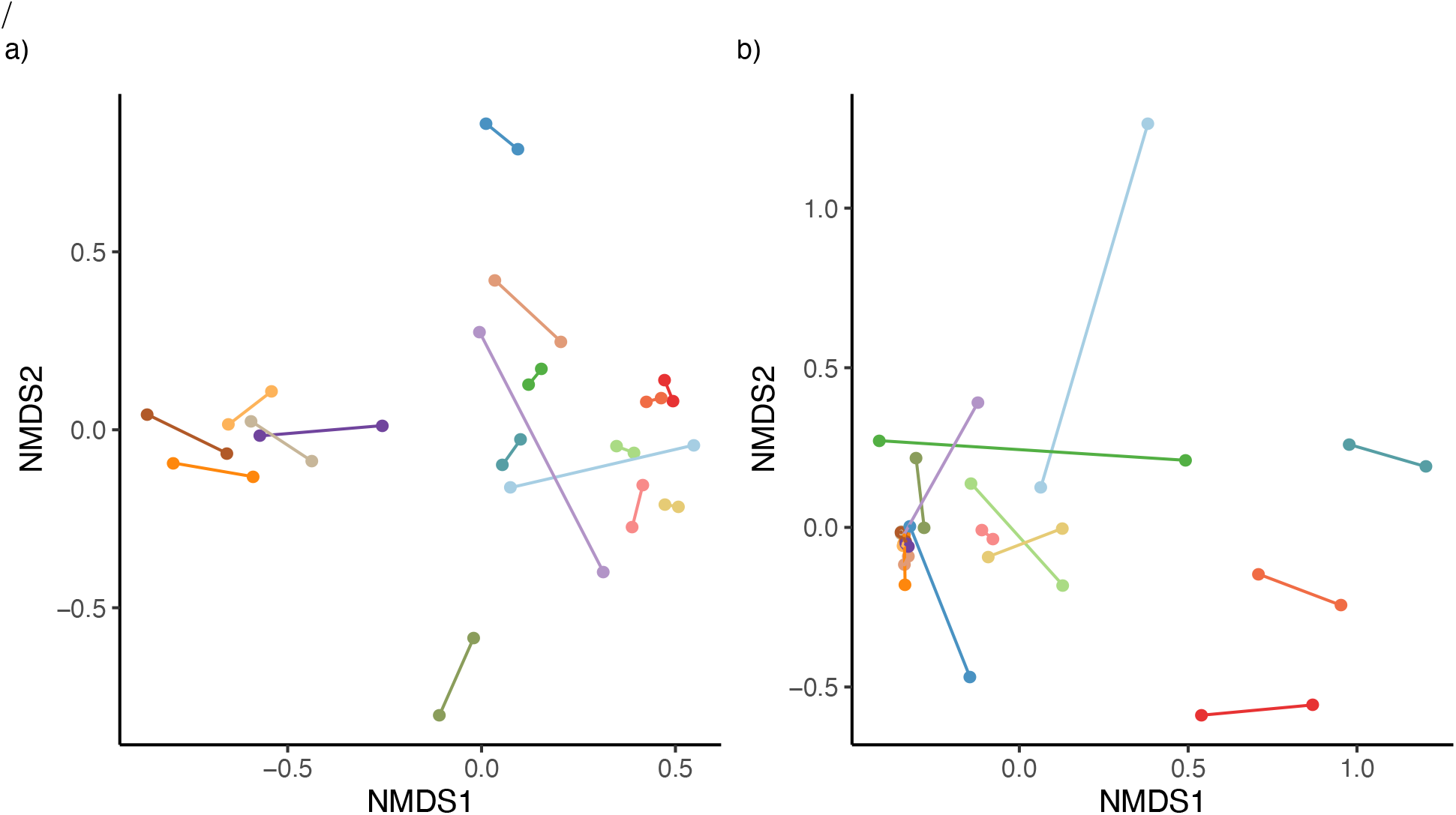
NMDS ordinations of seal louse microbiomes base on Bray–Curtis dissimilarity matrices. a) MAG matrix, and b) Kaiju matrix (species level). Lice originating from the same seal individual are colored similarly and connected by a line.

Similarly, in Kaiju matrices collapsed at the species level, most (>80% in all cases) of the variance was also explained by infrapopulation identity (PERMANOVA: Bray–Curtis, R^2^=0.804, F=4.346, P=0.001, Fig 3b; Jaccard, R^2^=0.803, F=4.319, P=0.001; Fig S3b). Again, results from simulations were similar [PERMANOVA: Bray-Curtis, R^2^ (min= 0.62, max= 0.88, mean= 0.75); P (min= 0.003, max= 0.058, n<0.05= 9/10); Jaccard, R^2^ (min= 0.63, max= 0.95, mean= 0.76), P (min= 0.002, max= 0.09, n<0.05= 9/10)]. Of all the others factors examined, only host status explained a significant amount of variance [PERMANOVA: Bray-Curtis, Host-status: R^2^= 0.22, F= 9.03, P= 0.001, Louse sex: R^2^= 0.08, F= 0.81, P= 0.564, Sequencing lane: R^2^= 0.01, F= 0.35, P= 0.859; Jaccard, Host-status: R^2^= 0.21, F= 8.73, P= 0.001, Louse sex: R^2^= 0.08, F= 0.88, P= 0.497, Sequencing lane: R^2^= 0.01, F= 0.4, P= 0.825; Mantel test, locality, Bray-Curtis: ρ (min= 0.04, max= 0.04, mean= 0.04), P (min= 0.5637, max= 0.5785, n<0.05= 0/10); Jaccard: ρ (min= −0.03, max= −0.03, mean= −0.03), P (min= 0.5337, max= 0.5488, n<0.05= 0/10)]. PERMANOVA analysis accounting for host status did not alter the significance of host identity (PERMANOVA: Bray-Curtis, R^2^= 0.52, F= 1.78, P= 0.007; Jaccard, R^2^= 0.59, F= 3.37, P= 0.001).

Furthermore, results were consistent when using matrices collapsed at the genus level (>77% of variance explained in all cases) (PERMANOVA: Bray–Curtis, R^2^= 0.865, F= 6.804, P= 0.001, Fig S4a; Jaccard, R^2^= 0.774, F= 3.634, P= 0.001; Fig S4b). Once again, results from simulations were similar [PERMANOVA: Bray-Curtis, R^2^ (min= 0.68, max= 0.96, mean= 0.8); P (min= 0.002, max= 0.073, n<0.05= 9/10); Jaccard, R^2^ (min= 0.54, max= 0.86, mean= 0.73), P (min= 0.003, max= 0.061, n<0.05= 9/10)]. Additionally, of all the others factors examined, only host status explained a significant amount of variance [PERMANOVA: Bray-Curtis, Host-status: R^2^= 0.3, F= 14, P= 0.001, Louse sex: R^2^= 0.05, F= 0.51, P= 0.851, Sequencing lane: R^2^= 0.01, F= 0.39, P= 0.753; Jaccard, Host-status: R^2^= 0.18, F= 7.19, P= 0.002, Louse sex: R^2^= 0.07, F= 0.75, P= 0.53, Sequencing lane: R^2^= 0.01, F= 0.40, P= 0.75; Mantel test, locality, Bray-Curtis: ρ (min= 0.09, max= 0.09, mean= 0.09), P (min= 0.7198, max= 0.7344, n<0.05= 0/10); Jaccard: ρ (min= 0.02, max= 0.02, mean= 0.02), P (min= 0.4043, max= 0.4246, n<0.05= 0/10)]. Likewise, PERMANOVA analysis accounting for host status did not alter the significance of host identity (PERMANOVA: Bray-Curtis, R^2^= 0.56, F= 4.73, P= 0.001; Jaccard, R^2^= 0.59, F= 2.96, P= 0.001).

## Discussion

Two different metagenomic approaches support a major role of infrapopulation identity (ringed seal host individual) in explaining microbiome variation among individuals of the seal louse. In addition, highly similar results were found for approaches using either presence-absence or quantitative matrices, suggesting that not only is bacterial composition, but also bacterial abundance explained by infrapopulation identity. Our analyses were done on whole louse individuals and, thus, we cannot confidently differentiate between bacterial taxa inhabiting the lice (e.g., *Wolbachia* or *Hodgkinia*) from transient taxa present in the host blood meal (e.g., *Chlamydia*). Nevertheless, in line with current evidence on the determinants of microbiome composition of bloodsucking parasites, the louse blood meal from individual seals is the most likely candidate in explaining the patterns of microbiome variation across the louse infrapopulations found here. Indeed, many of the taxa found in our analyses have already been found in other bloodsucking parasites, thus supporting the influence of blood in shaping the composition of parasite microbiomes studied here (Jiménez-Cortés et al. 2018).

However, other factors in addition to blood may have contributed to the similarity of microbiomes between individual lice from the same seal host individual. Some similarity may have arisen from shared environmental bacteria, those on the surface of the louse from a shared environment (skin and fur of the host), or contamination between louse individuals in screw-cap tubes, and not filtered by our decontamination procedures. There may also be insect-specific bacterial taxa, independent from the host blood, that are shared horizontally between individual lice from the same infrapopulation. Finally, louse infrapopulations are known to typically be highly inbred, with a high level of relatedness between individuals (Koop et al. 2014, DiBlasi et al 2018, Virrueta Herrera et al., in prep.). It may be that there are louse genetic factors that interact with the microbiome to produce a specific composition (Blekhman et al. 2015; Dobson et al. 2015; Suzuki et al. 2019).

Our results are congruent with previous findings on the influence of host blood on microbiomes of bloodsucking parasites. Specifically, several studies have found a major role of the specific host species from which a blood meal is taken in shaping microbiomes of other bloodsucking organisms, such as ticks (*Ixodes scapularis, Ixodes pacificus*) and mosquitoes (*Aedes aegypti*) (Swei and Kwan 2017; Landesman et al. 2019; Muturi et al. 2019). Furthermore, Landesman et al. (2019) showed that microbiomes of deer tick (*Ixodes scapularis*) nymphs were largely explained by the individual hosts of the tick, a result similar to the one obtained here. Interestingly, in that study, the percentage of variation explained was considerably lower (45%) than that found here (>77%). It may be that differences in parasite ecology, such as the whether the parasite is a permanent or a recurrent feeder (as are both the case in sucking lice) may modulate the extent to which host blood shapes parasite microbiomes. The differences in the proportion of variance explained by infrapopulation identity between the two studies could also be due to differences in experimental design, such as the number of sampled infrapopulations (3 in ticks, and 17 in the seal lice here) and whether the sample design is balanced (i.e., the same number of individual parasites sampled per infrapopulation).

The knowledge that blood from the same individual seal host may influence the similarity of the microbiome of blood-feeding lice from that host can potentially provide new insights into the influence of host blood on such parasites. There are least two not necessarily mutually exclusive processes may explain the influence of a host individual’s blood on louse microbiomes. First, the blood from a particular host individual may contain a specific composition of bacterial loads that enter the louse on consumption of blood. Indeed, anopluran lice might have a higher likelihood of being colonized by bacteria from host blood because they do not possess a peritrophic membrane, an extracellular layer in the midgut that is composed of chitin, proteoglycans, and proteins, which in most other insects surrounds the ingested food bolus and separates the gut content, including bacteria, from the epithelium (Terra 2001; Waniek 2009). Indeed, the idea that a lack of a peritrophic membrane may facilitate colonization of blood-feeding parasites by bacteria present in the host blood has potentially also been supported by work on mouse fleas (*Rhadinopsylla dahurica*), which also lack this membrane (Li et al. 2018). In this case, there was evidence of homogenization (i.e., similar bacterial communities) between the host blood and the parasite (whole flea individuals). The lack of a peritrophic membrane is often associated with permanent parasites, such as blood-feeding lice, for which the continual availability of food means that there is less selection for efficiency of digestion. Therefore, the presence versus absence of a peritrophic membrane may explain the differences between lice and ticks (of which the latter possess a peritrophic membrane) with regards to the influence of host blood on the composition of the parasite microbiome.

A second possibility that could explain why host blood may influence louse microbiome composition is that the conditions during blood digestion may alter bacterial taxa that were already present in the louse. The specifics of blood digestion may have an individual host-specific signature. Specifically, catabolism of blood meal leads to the generation of reactive oxygen species that are known to alter the midgut bacterial composition and diversity of bloodsucking parasites (Souza et al. 1997; Wang et al. 2011; Muturi et al. 2019). Also, the blood meals of different host species are also known to differ in composition (e.g., total protein, hemoglobin, and hematocrit content), and these differences may lead to a differential proliferation of microbial taxa during digestion by the parasite (Souza et al. 1997; Wang et al. 2011; Muturi et al. 2019). It may be the case that differences in blood composition among individuals even within the same host species may be shaping the bacterial composition of lice in a manner that is specific to host individuals.

Bloodsucking organisms, and anopluran lice in particular, are well known to rely on mutualistic endosymbionts to complement deficiencies in their diet (Perotti et al. 2008; Boyd and Reed 2012; Boyd et al. 2017; Jiménez-Cortés et al. 2018). Notwithstanding that several of the bacterial taxa we found may not be stable inhabitants of lice, we did find evidence for the presence of several louse-specific bacterial taxa. These include the obligate intracellular arthropod bacteria *Wolbachia* (Werren 1997) and *Hodgkinia* (for which only endosymbionts of *Cicadas* are known; McCutcheon et al. 2009). Accordingly, we explored our MAGs for genome characteristics typical of endosymbionts. In particular, because endosymbiont genomes typically are small and have an AT bias, we explored the relative position of the observed MAGs in a “Genome size ~ GC content” correlation plot (Wernegreen 2015; Figure 4). Bin 1 appears to be the best candidate to be a mutualistic endosymbiont, according to its relative position in the correlation plot. This MAG was 100% complete (according to CheckM; Parks et al. 2015), detected in most samples (prevalence = 71%), and classified with confidence as Flavobacteriaceae. MiGA analyses suggest it may even belong to *Chryseobacterium* (p-value 0.585). Endosymbionts belonging to *Chryseobacterium* are known in other arthropods (e.g., termites, mosquitoes, cockroaches, and ticks; Eutick et al. 1978; Dugas et al. 2001; Campbell et al. 2004; Montasser 2005; Burešová et al. 2006). Additionally, we conducted a preliminary investigation of the metabolic capabilities of this bacterium by investigating the completeness of metabolic pathways using GhostKOALA (Kanehisa et al. 2016) and KEGG-Decoder (Graham et al. 2018). This MAG has complete routes for synthesis of vitamin B (riboflavin), an essential amino acid (lysine), and several non-essential amino acids (e.g., serine; see Table S4), as well as many fully or partially missing routes that may be redundant or potentially shared (or synthesized along) with the louse (Table S4).

**Figure 4.**
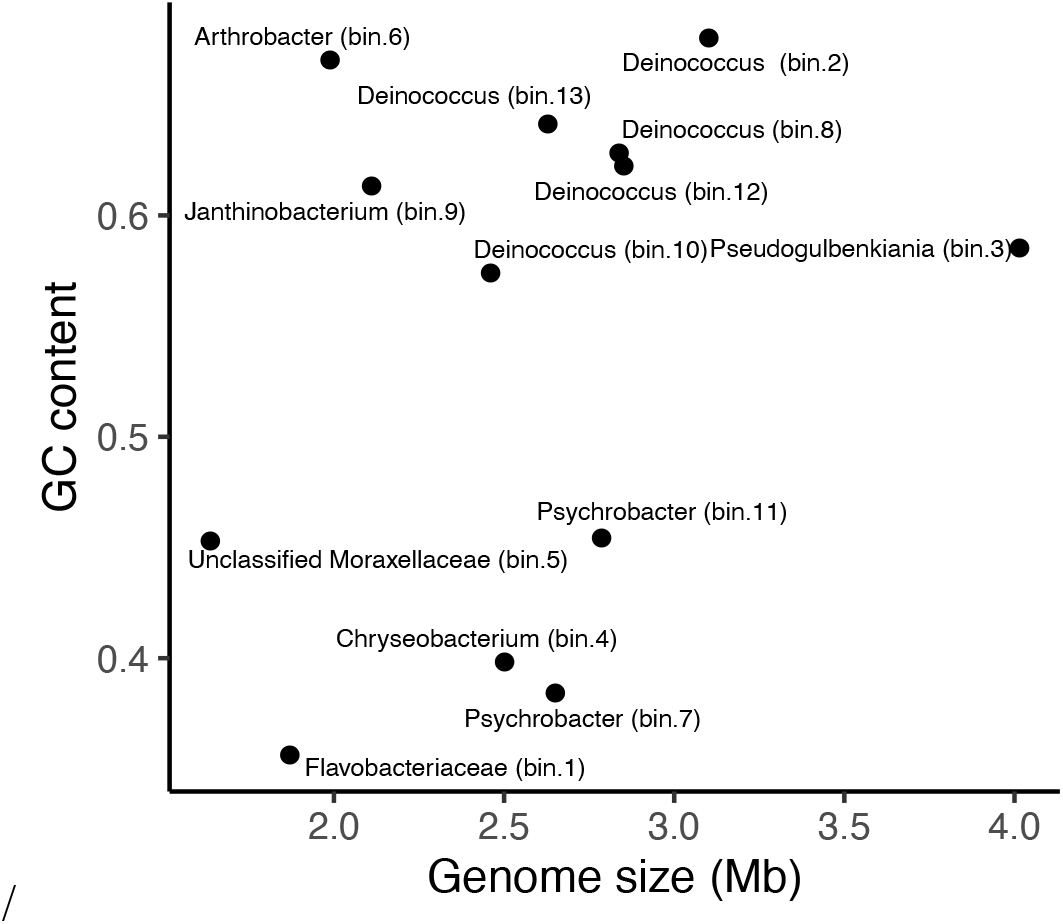
Scatter plot showing the relationship between genome size (Mb) and GC content (i.e., proportion of G and C sites) for sequenced MAGs.

Overall, these results are congruent with what has been found for endosymbionts of bloodsucking parasites (Moriyama et al. 2015; Boyd et al. 2016; Santos-Garcia et al. 2017; Duron et al. 2018). Another anopluran pinniped louse (*Proechinophthirus fluctus*) has been found to have a *Sodalis* endosymbiont (Boyd et al. 2016), but we found no evidence of *Sodalis* in *Echinophthirius horridus*. Other species of Anoplura have yet other endosymbionts (Boyd et al. 2017, Ríhová et al. 2017), suggesting that endosymbiont replacement is an ongoing and relatively common process within the order. Further research, including phylogenomic studies improving the phylogenetic placement of this potentially mutualistic bacterium and studies using fluorescence in situ hybridization (FISH) to ascertain the location of this bacterium in louse individuals, is needed to get deeper insight into the interaction of this bacterium with *E. horridus*.

Finally, at a broader scale, our results are congruent with previous studies that have found a major role of different levels of subdivision in shaping microbiomes in a wide range of systems. For instance, population identity has been found to largely explain microbiome composition of great apes (Campbell et al. 2020), American pikas (*Ochotona princeps*) (Kohl et al. 2018), and humans (Rothschild et al. 2018). Similar results have also been found across subdivision levels other than populations, such as ecotypes (human lice *Pediculus humanus*; Agany et al. 2020), or by spatial proximity (North American moose *Alces alces;* Fountain-Jones et al. 2020). On the other hand, while the influence of subdivision on microbiome composition is widely supported, much less is known about whether signs of phylosymbiosis can be found at these levels. Kohl et al. (2018) found that populations largely explained microbiome variation in American pikas along with a phylosymbiotic pattern (i.e., closely related hosts had similar microbial communities). Interestingly, while we did not investigate the presence of phylosymbiosis here, our results suggest that a phylosymbiotic pattern is not always found at an intraspecific level. For instance, this may be the case of subdivided systems in which the food source (which allows the dispersal of microbes from one organism to another) constitutes the main determinant of microbiome composition, and populations are not necessarily composed of closely related organisms. However, phylosymbiotic patterns across species have been found in relatively similar systems. Therefore, bacterial dispersal, ecological drift, diversification, or microbe-microbe interactions may be the main factors explaining phylosymbiosis in these systems. More studies on the origin and prevalence of phylosymbiotic patterns within and across species are clearly needed.

## Acknowledgments

We thank all researchers and students for collecting the lice analyzed in this study. Especially, we would like to thank researchers Miina Auttila, Vincent Biard, Meeri Koivuniemi, Lauri Liukkonen, Marja Niemi, Sari Oksanen, Mia Valtonen, and Eeva Ylinen for keeping eye on the lice during their own studies.

## Author contributions

J.D., S.V.H, and K.P.J. conceived the study. T.N. and M.K. obtained samples. S.V.H. and K.P.J. collected the data. J.D. analyzed the data. T.N., M.V., and K.P.J. obtained financial support for the project. J.D. wrote the manuscript and all authors contributed to editing the manuscript.

## Funding

This study was supported by the US National Science Foundation (DEB-1239788, DEB-1342604, and DEB-1926919 to K.P.J). Sequencing costs were supported by grants from the Oskar Öflund Foundation, the Betty Väänänen Foundation, Societas Pro Fauna et Flora Fennica, and the Nestori Foundation.

## Availability of data and material

Raw sequence reads for all samples are available under (NCBI) project *ACCESSION CODE*. Metagenomic assemblies are available at Figshare (doi: 10.6084/m9.figshare.12366575 —private link for review: https://figshare.com/s/171207f7fb73f7d290a6). Metagenomic assemblies are also available from NCBI Genome (*ACCESION CODES*).

## Ethics approval and consent to participate

Telemetry studies have been approved by the local environmental authority Centre for Economic Development, Transport and the Environment (permit numbers: ESAELY/433/ 07.01/2012 and ESA-2008-L-519-254) and the Animal Experiment Board in Finland (permit numbers: ESAVI/ 8269/04.10.07/2013 and ESAVI-2010-08380/Ym-23).

## Competing interests

The authors declare that they have no competing interests.

## Notes

### Competing Interest Statement

The authors have declared no competing interest.

### Summary of Updates

We have modified the acknowledgment section (as suggested by a reviewer) and corrected some typos.

