## Supplementary figures and images for "Patterns of microbiome variation among infrapopulations of permanent bloodsucking parasites"

### Figure S1

a) MAGs matrix

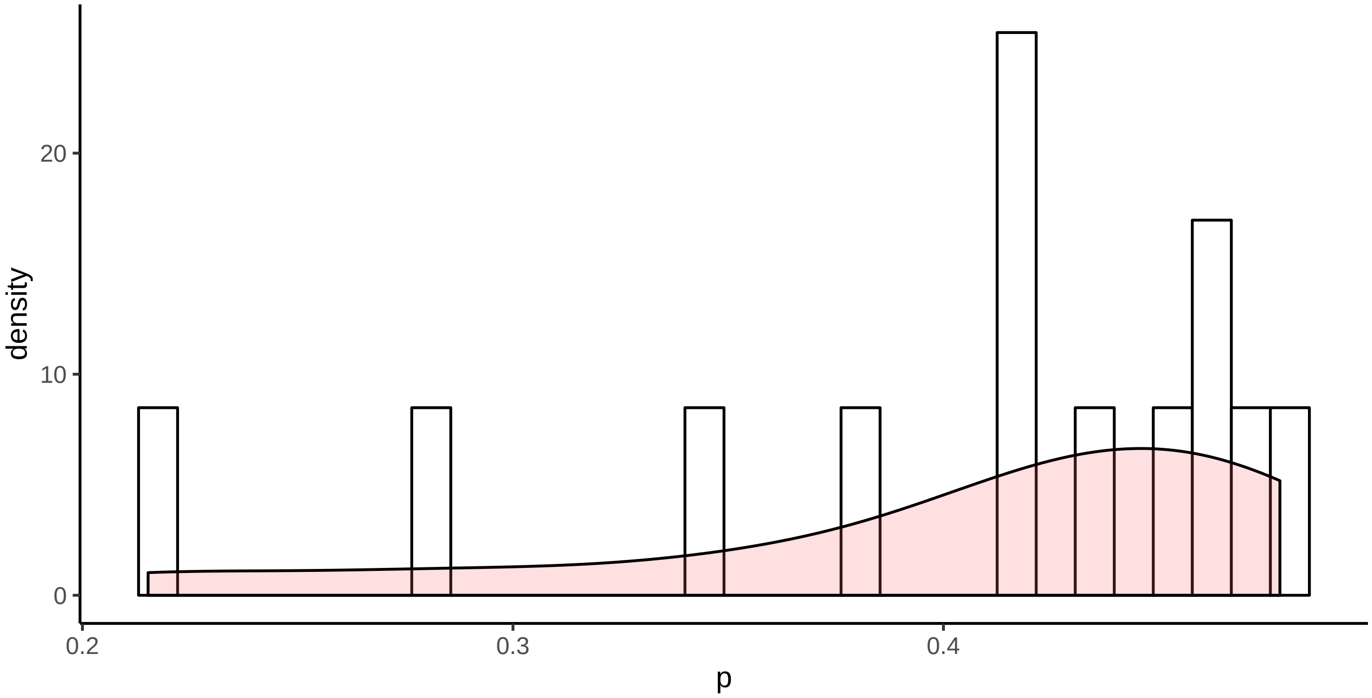


b) Kaiju matrix (species level)

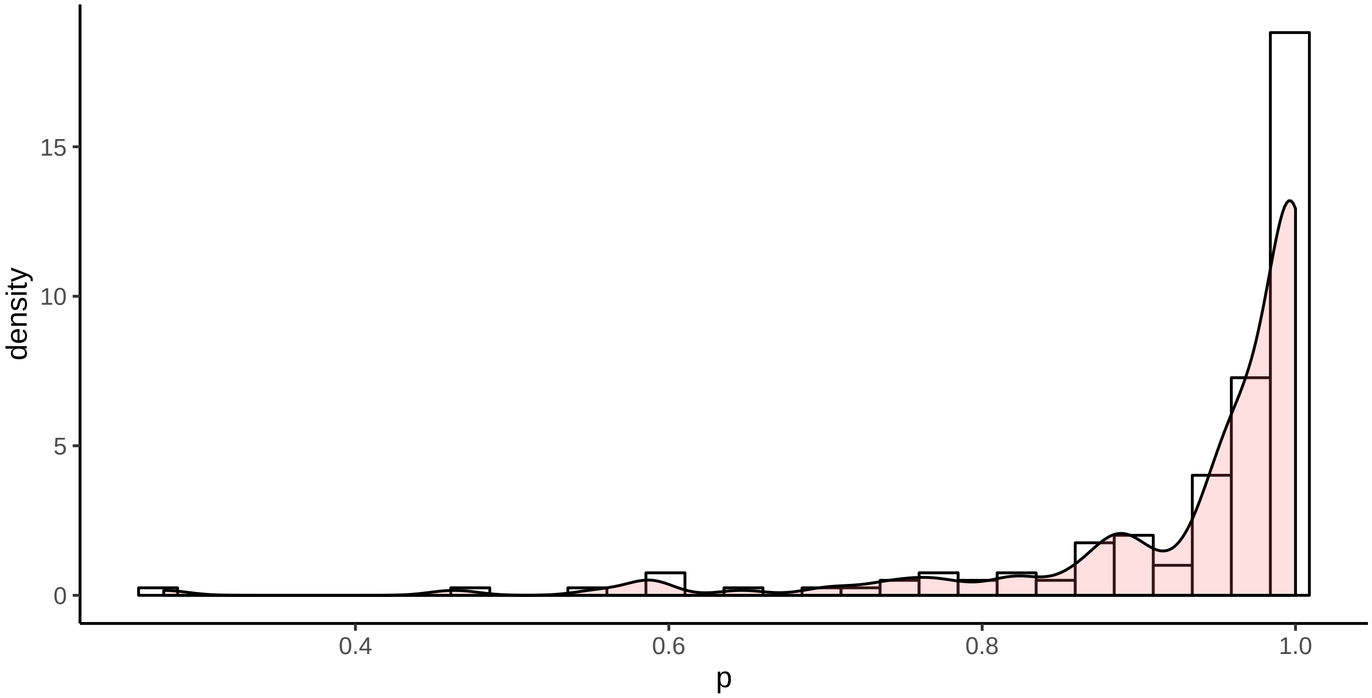


c) Kaiju matrix (genus level)

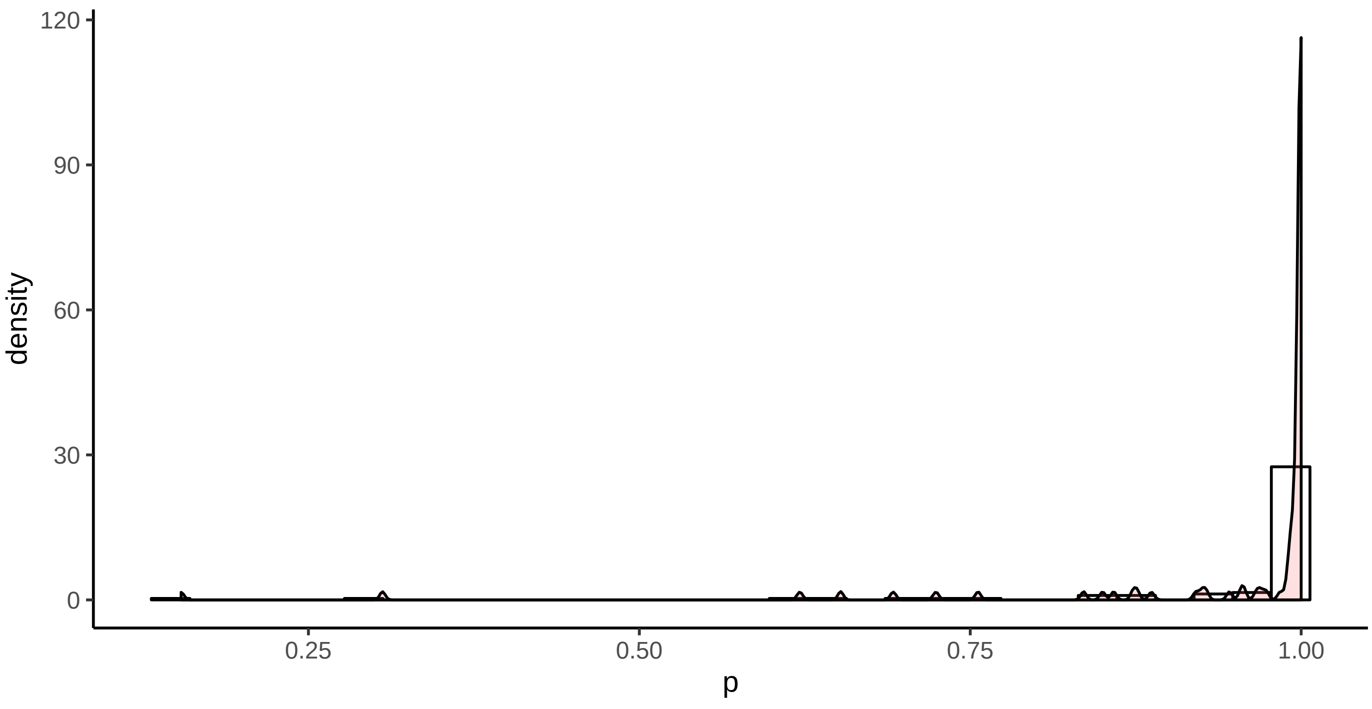

### Figure S2

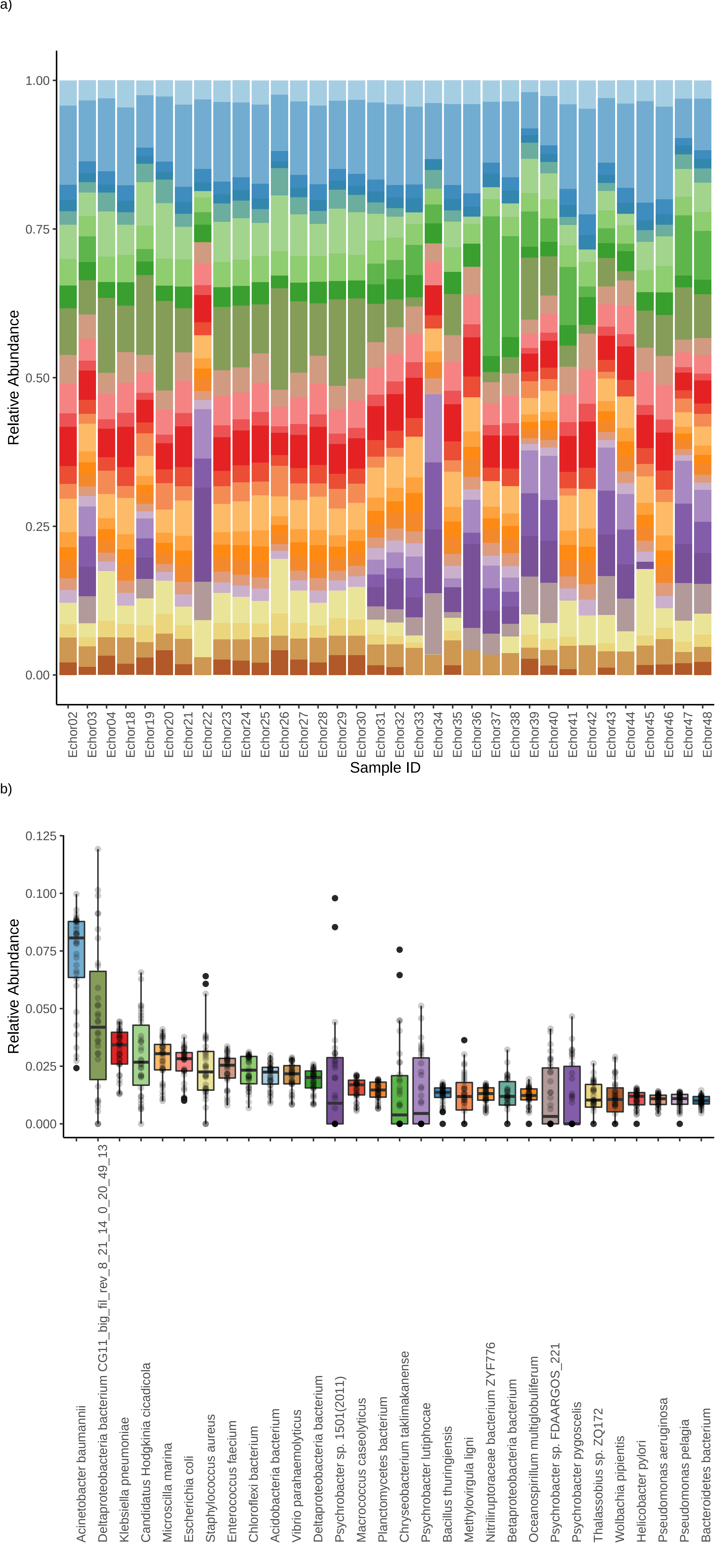

### Figure S3

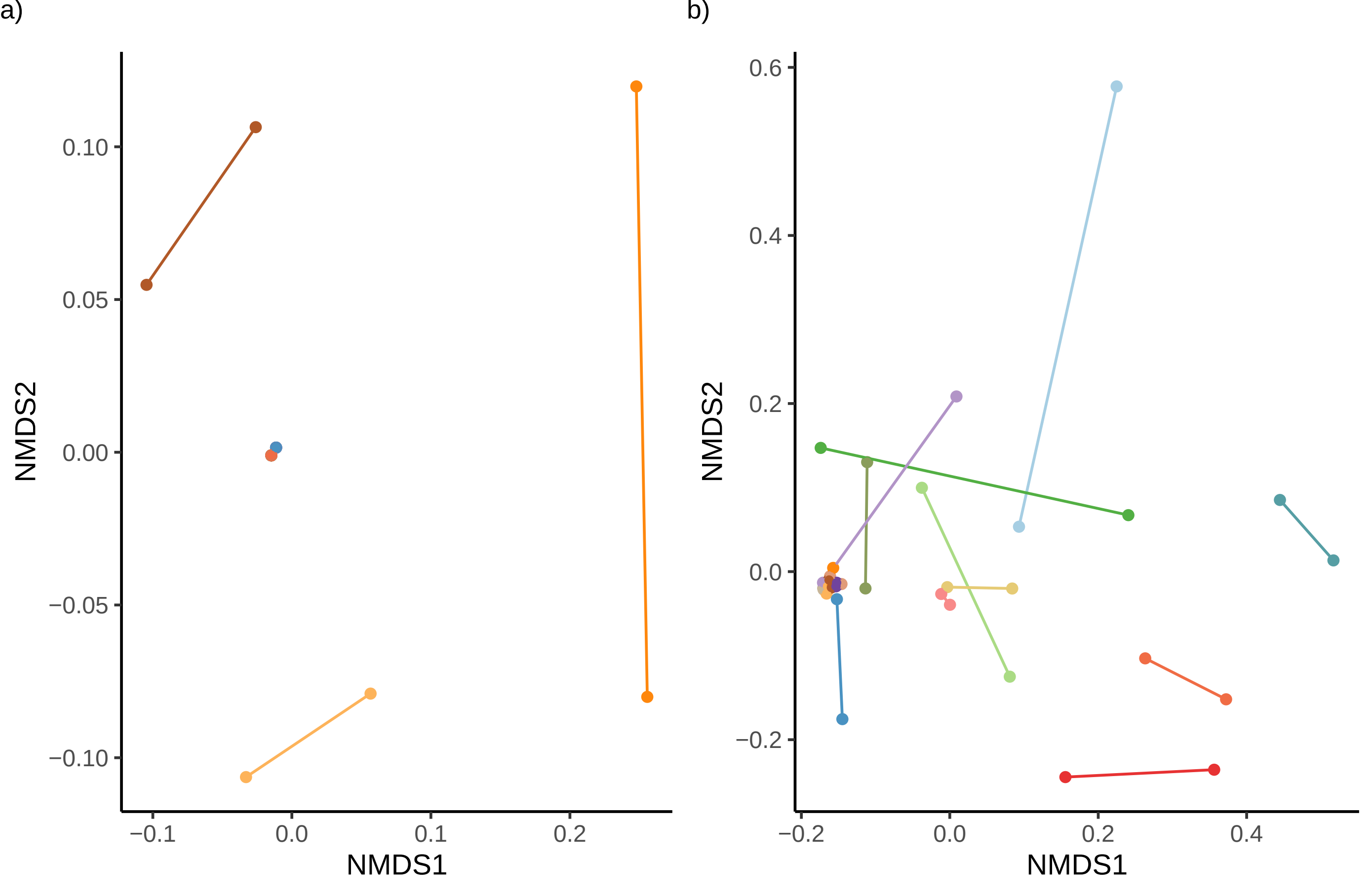

### Figure S4

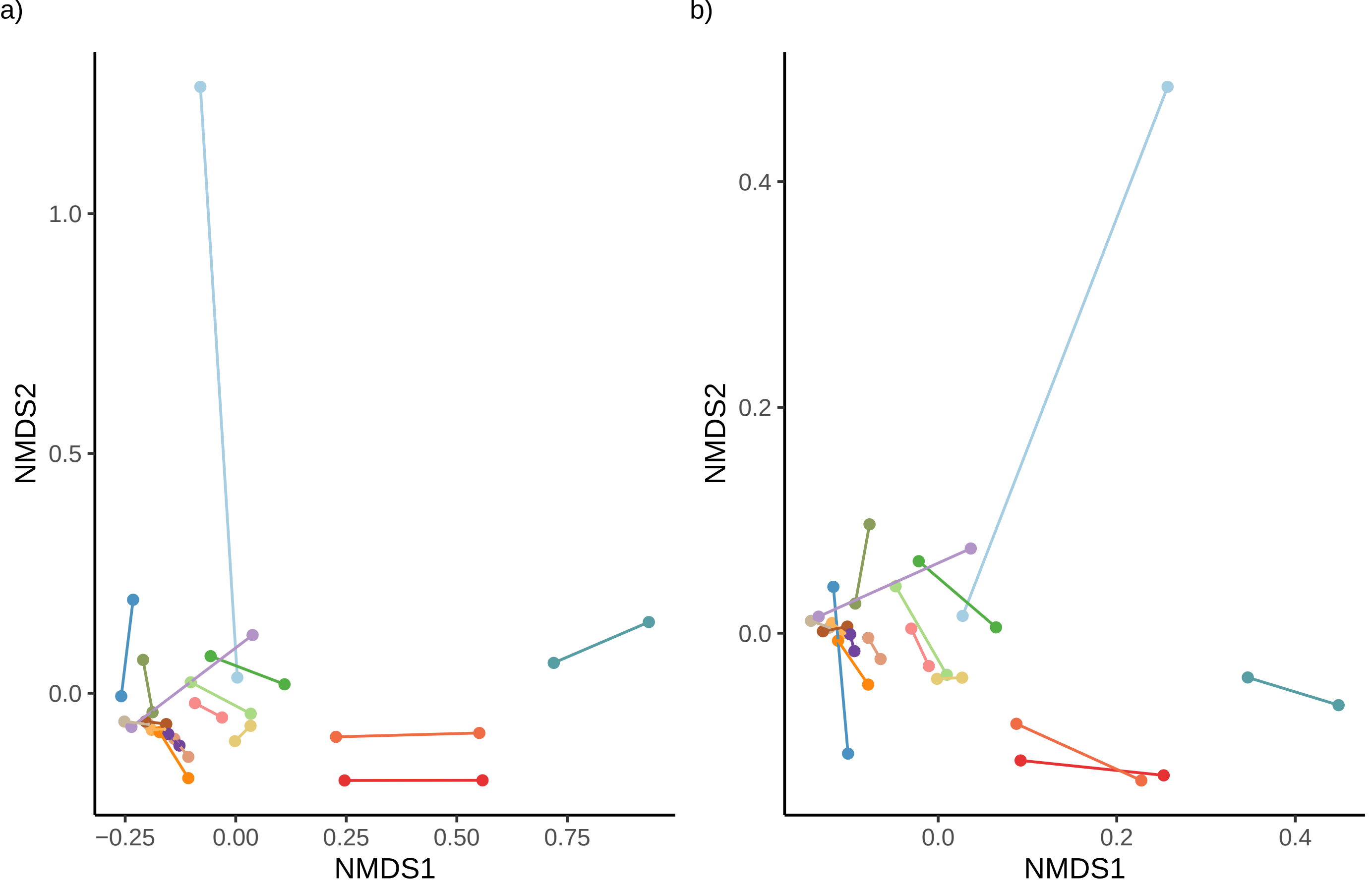
